# *dCBP*-mediated histone lactylation contributes to meiotic chromosome maintenance

**DOI:** 10.64898/2026.05.15.725312

**Authors:** Keisuke Nakayama, Daisuke Saito, Yoshiki Hayashi

## Abstract

Histone lactylation is a recently identified histone post-translational modification (PTM) that links energy metabolism to chromatin regulation. Although histone lactylation has been implicated in transcriptional activation, its function in meiotic chromatin remains unclear. Previously, we identified enrichment of multiple histone lactylation marks within the meiotic karyosome, a highly condensed and transcriptionally repressive chromatin structure formed in *Drosophila* oocytes. Here, through an RNAi-based screen, we identified the CBP family protein *dCBP* as a regulator of histone lactylation in the karyosome. Germline-specific knockdown of *dCBP* preferentially reduced histone lactylation, including H4K8 lactylation, and caused premature disruption of the synaptonemal complex, abnormal egg chamber development with excess nurse cells, reduced egg production, and decreased embryonic viability. Corresponding histone acetylation marks were comparatively less affected than histone lactylation by *dCBP* knockdown. Together, our findings provide evidence that *dCBP*-mediated histone lactylation contributes to meiotic chromosome maintenance and suggest a potential link between energy metabolism and meiotic chromatin regulation.

## Introduction

Histone lysine post-translational modifications (PTMs) are central regulators of chromatin architecture and gene expression. Among them, histone lactylation, a recently identified PTM derived from glycolytic lactate, has emerged as a potential link between energy metabolism and epigenetic regulation (Zhang et al., 2019). Although histone lactylation has been implicated in transcriptional activation in several biological contexts, including tumorigenesis, inflammation, and neurological diseases (Peng and Du, 2025, Hu et al., 2024, Li et al., 2024, Zhang et al., 2019), whether histone lactylation contributes to specialized chromatin states requiring extensive chromosome reorganization, such as those occurring during meiosis, remains unknown.

Meiosis is a specialized cell division program that generates haploid gametes and promotes genetic diversity. During meiosis, chromosomes undergo extensive reorganization, including homolog pairing, synapsis, meiotic recombination, and large-scale chromatin remodeling. Consistent with a potential role for histone lactylation in meiotic chromatin regulation, we recently found that histone lactylation is strongly enriched on meiotic chromosomes in both *Drosophila* and mouse oocytes (Hayashi et al., 2025c).

In the *Drosophila* ovary, germline stem cell-derived cysts give rise to a single oocyte and 15 nurse cells through incomplete cytokinesis. During cyst development, meiotic chromosome structures such as the synaptonemal complex (SC) initially form in multiple cyst cells, but meiotic progression subsequently becomes restricted to the oocyte (Hughes et al., 2018). Following egg chamber formation, oocyte chromatin undergoes dramatic compaction into a specialized configuration of meiotic chromosomes termed the karyosome, whereas nurse cells instead become highly polyploid and transcriptionally active. Notably, histone lactylation is strongly enriched in the karyosome (Hayashi et al., 2025c).

Although the striking enrichment of histone lactylation in the karyosome suggests a potential role in meiotic chromosome regulation, the mechanisms regulating histone lactylation in oocytes remain poorly understood. Here, through a functional knockdown screen, we identified the *Drosophila* orthologue of CREB-binding protein (*dCBP*), *nejire*, as a critical regulator of histone lactylation in the karyosome. Notably, CBP/p300 family proteins have previously been implicated in histone lactylation in mammalian cells (Zhang et al., 2019, Li et al., 2024). Interestingly, germline-specific inhibition of *dCBP* caused defects in SC maintenance and oocyte specification. Together, our findings identify *dCBP* as a key regulator of histone lactylation in the oocyte and suggest that histone lactylation contributes to meiotic chromosome regulation.

## Results and Discussion

### Identification of a factor regulating lactylation in the karyosome

To investigate the molecular mechanisms regulating histone lactylation in the karyosome, we performed an RNA interference (RNAi)-mediated knockdown screen of candidate genes in the germline using the germline-specific Gal4 driver, *nanos* (*nos*)-Gal4 (hereafter, we describe the genotype of germline-specific RNAi of genes as “*nos* > gene RNAi”). We selected seven candidate genes previously implicated in histone lactylation or related acylation pathways in mammals, including metabolic enzymes and histone acetyltransferases (Figure 1B, Table 1). These candidate genes included *Ldh* (CG10160) (Zhang et al., 2019), *AcCoAS* (CG9390) (Zhu et al., 2025), CG6432 (the *Drosophila* orthologue of human ACCS3)(Zhu et al., 2025), *Gcn5* (CG4107)(Zhu et al., 2025), *dCBP* (CG15319) (Zhang et al., 2019), *Tip60* (CG6121) (Chen et al., 2024), and *AlaRS-m* (CG4633) (Mao et al., 2024). We used a pan-lysine lactylation antibody (pan-KLa) to assess lactylation signals in the karyosome. To confirm germline expression of these candidates, we examined a published scRNA-seq dataset and found that most genes were expressed in the ovarian germline, except for CG6432 (Supp. Figure 1). Among the candidates, knockdown of *dCBP*, a known transcriptional co-activator and histone acetyltransferase (HAT)(Feller et al., 2015), was the only condition that showed a significant reduction in karyosome lactylation (Figures 1A-C, Supp. Table 1). These results identified *dCBP* as a candidate regulator of histone lactylation in the karyosome, prompting further investigation into its role in karyosome regulation during oogenesis.

**Figure 1.**
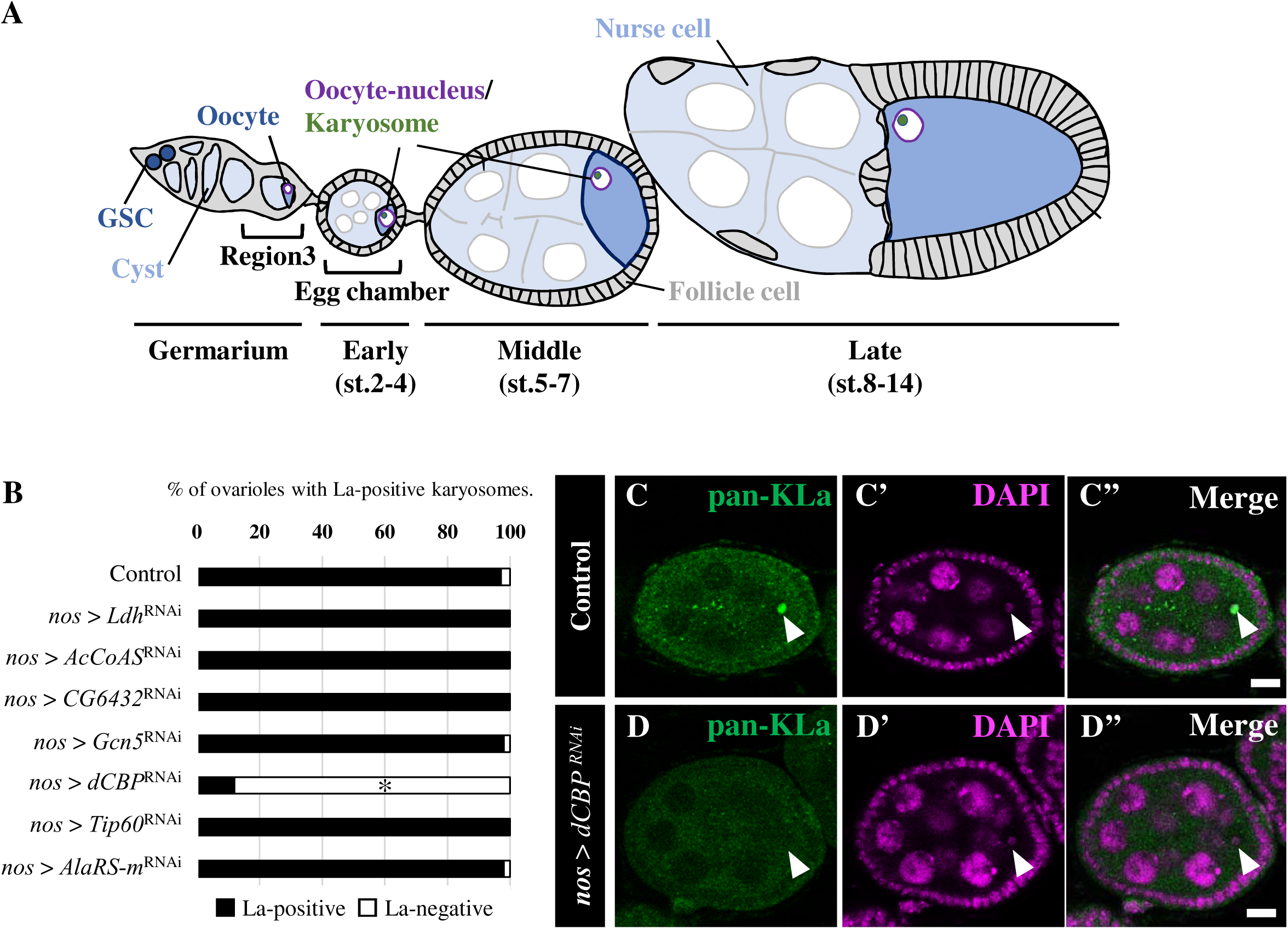
*dCBP* is required for lysine lactylation in the karyosome. (A) Schematic diagram of *Drosophila* oogenesis. Germline stem cells (GSCs) are located at the anterior tips of germarium. GSC-derived germline cysts give rise to a single oocyte and 15 nurse cells through incomplete cytokinesis within the germarium. Formed germline cysts are encapsulated with surrounding somatic follicle cells to form egg chambers (Hayashi et al., 2020). Within the cysts, oocyte nuclei maintain the meiotic chromosome; conversely, nurse cell nuclei become high-ploidy with endocycle. In this study, stages 2–4, 5–7, and 8 and later are defined as early, mid-, and late oogenesis, respectively. (B) RNAi-mediated knockdown screen to identify factors that regulate lysine lactylation in the karyosome. (C–D’’) Lysine lactylation in the karyosome of control (C–C’’) and germline-specific *dCBP*-knockdown ovaries (D–D’’). Green indicates lysine lactylation, and magenta indicates DAPI staining of DNA. Arrowheads indicate karyosomes. Scale bars: 10 µm. Asterisks indicate p < 0.05 by Fischer’s exact test.

### dCBP regulates histone lysine lactylation in the karyosome

Because the pan-KLa antibody recognizes lactylated lysines broadly, including non-histone proteins, we next investigated whether *dCBP* regulates lactylation at specific histone lysine residues. We therefore examined the effects of *dCBP* knockdown on individual histone lactylation marks using antibodies recognizing specific histone lactylation marks. We chose several lactyl-lysine modifications on histone H3 and H4 [H3K14La (Figures 2A-A’’), H4K8La (Figures 2D-D’’), and H4K12La (Figures 2G-G’’)], which we previously found to be enriched in the karyosome (Hayashi et al., 2025c). Quantitative image analysis showed that all examined histone lactylation marks were significantly enriched in the karyosome in mid-oogenesis relative to surrounding follicle cells, with H4K8La showing the strongest enrichment (Supp. Figures 2A-C). Lactylation levels at these histone lysines were approximately four-fold (H3K14La) to twenty-fold (H4K8La) higher in the karyosome than in surrounding follicle cells, suggesting that these histone lactylation marks are highly enriched in the karyosome. We next examined whether these histone lactylation marks were affected by *dCBP* knockdown, and found that all examined histone lactylation marks were reduced to approximately one-third to one-fourth of control levels [H3K14La (Figures 2A-C), H4K8La (Figures 2D-F), H4K12La (Figures 2G-I)]. Thus, our results strongly suggest that *dCBP* plays a central role in histone lysine lactylation in the karyosome.

**Figure 2.**
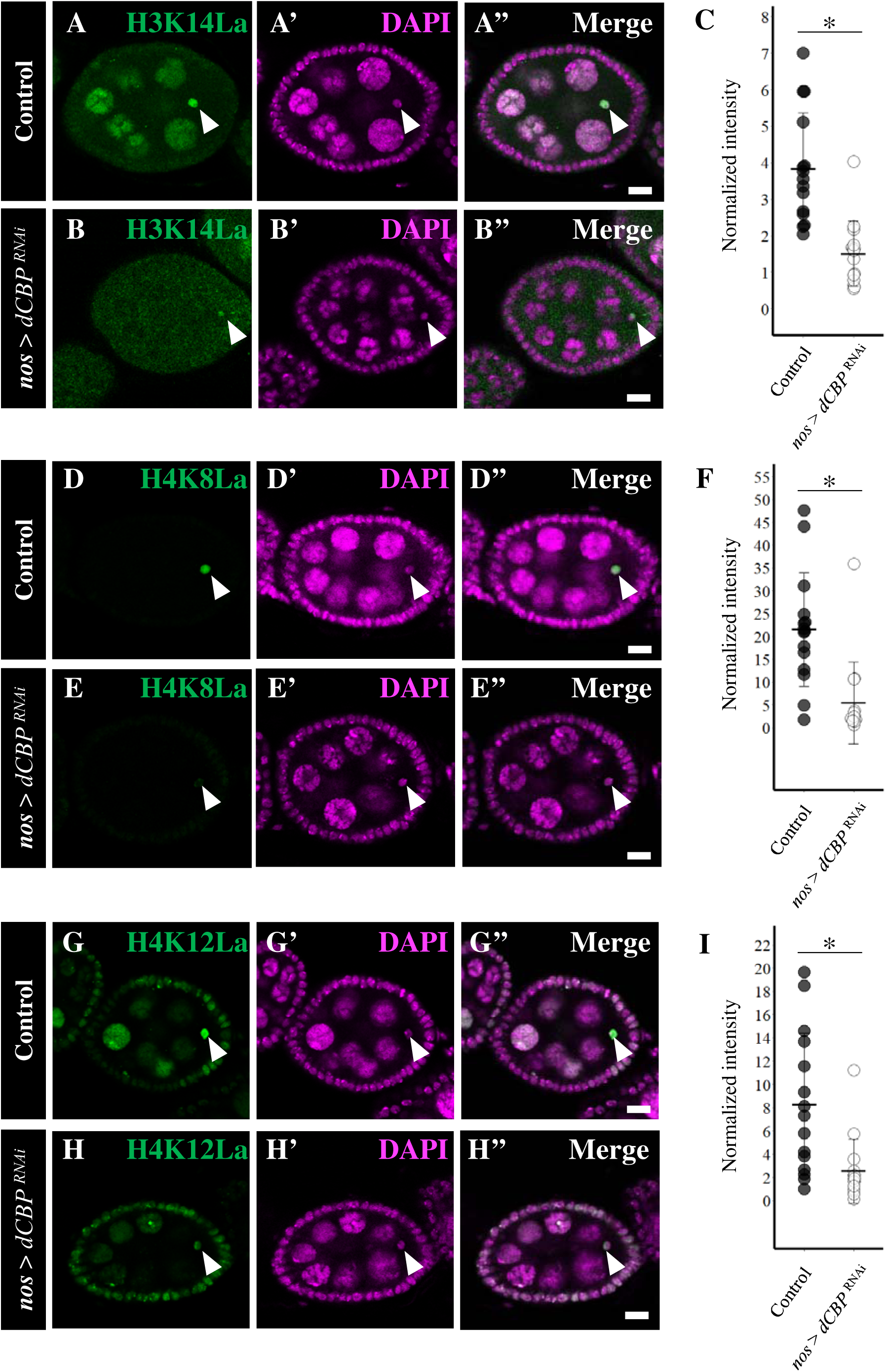
*dCBP* plays a central role in histone lysine lactylation in the karyosome. (A–C) H3K14 lactylation in control (A–A’’) and germline-specific *dCBP*-knockdown (B–B’’) karyosomes. Quantification of normalized karyosome signal intensity is shown in (C) (control, 3.83±1.52 S.D.; *nos* > *dCBP*^RNAi^, 1.51±0.89 S.D.; p <0.05, Welch’s t-test). (D–F) H4K8 lactylation in control (D–D’’) and germline-specific *dCBP*-knockdown (E–E’’) karyosomes. Quantification of normalized karyosome signal intensity is shown in (F) (control, 21.51±12.48 S.D.; *nos* > *dCBP*^RNAi^, 5.44±8.97 S.D.; p<0.05, Welch’s t-test). (G–I) H4K12 lactylation in control (G–G’’) and germline-specific *dCBP*-knockdown (H–H’’) karyosomes. Quantification of normalized karyosome signal intensity is shown in (I) (control, 8.30±6.09 S.D.; *nos* > *dCBP*^RNAi^, 2.55±2.75 S.D.; p<0.05, Welch’s t-test). For normalization methods, see Materials and Methods. In all images, green indicates histone lactylation and magenta indicates DAPI staining of DNA. Arrowheads indicate karyosomes. Scale bars: 10 µm. Asterisks indicate p < 0.05 (Welch’s t-test).

### Active chromatin-associated histone modifications accumulate in meiotic karyosome chromatin

Previous studies have shown that histone lactylation is associated with transcriptional activation (Zhang et al., 2019). Because the karyosome consists of highly compacted meiotic chromatin that is largely transcriptionally inactive, our findings suggest that histone modifications generally linked to active chromatin may also accumulate in this specialized chromatin environment. Consistent with this idea, a previous study reported the presence of histone acetylation within the karyosome (Navarro-Costa et al., 2016). Since dCBP is a well-characterized histone acetyltransferase in *Drosophila*, we next examined whether acetylation at the same lysine residues undergoing lactylation is also enriched in the karyosome. To address this question, we performed immunostaining using antibodies recognizing specific acetylated histone lysines, including H3K14Ac, H4K8Ac, and H4K12Ac. All examined acetylation marks were enriched in the karyosome relative to surrounding follicle cells; however, the degree of enrichment was substantially weaker than that observed for the corresponding lactylation marks (Supp. Figures 2D-F). These observations suggest that histone lactylation is more prominently associated with these lysine residues in karyosome chromatin than histone acetylation.

We next investigated whether *dCBP* contributes to acetylation at these sites within the karyosome. Although *dCBP* knockdown significantly reduced H4K12Ac levels (Figures 3G-I), the effects on H3K14Ac and H4K8Ac were comparatively modest (Figures 3A-F). Notably, H4K8Ac showed the weakest sensitivity to *dCBP* knockdown, whereas H4K8La displayed the strongest karyosome enrichment among the lactylation marks examined (Figure 2F; Supp. Figure 2B). Together, these findings suggest that although histone acetylation is present in transcriptionally inactive karyosome chromatin, histone lactylation may represent the predominant acyl modification at these residues. Our results further support the idea that *dCBP* preferentially contributes to histone lactylation in the karyosome.

**Figure 3.**
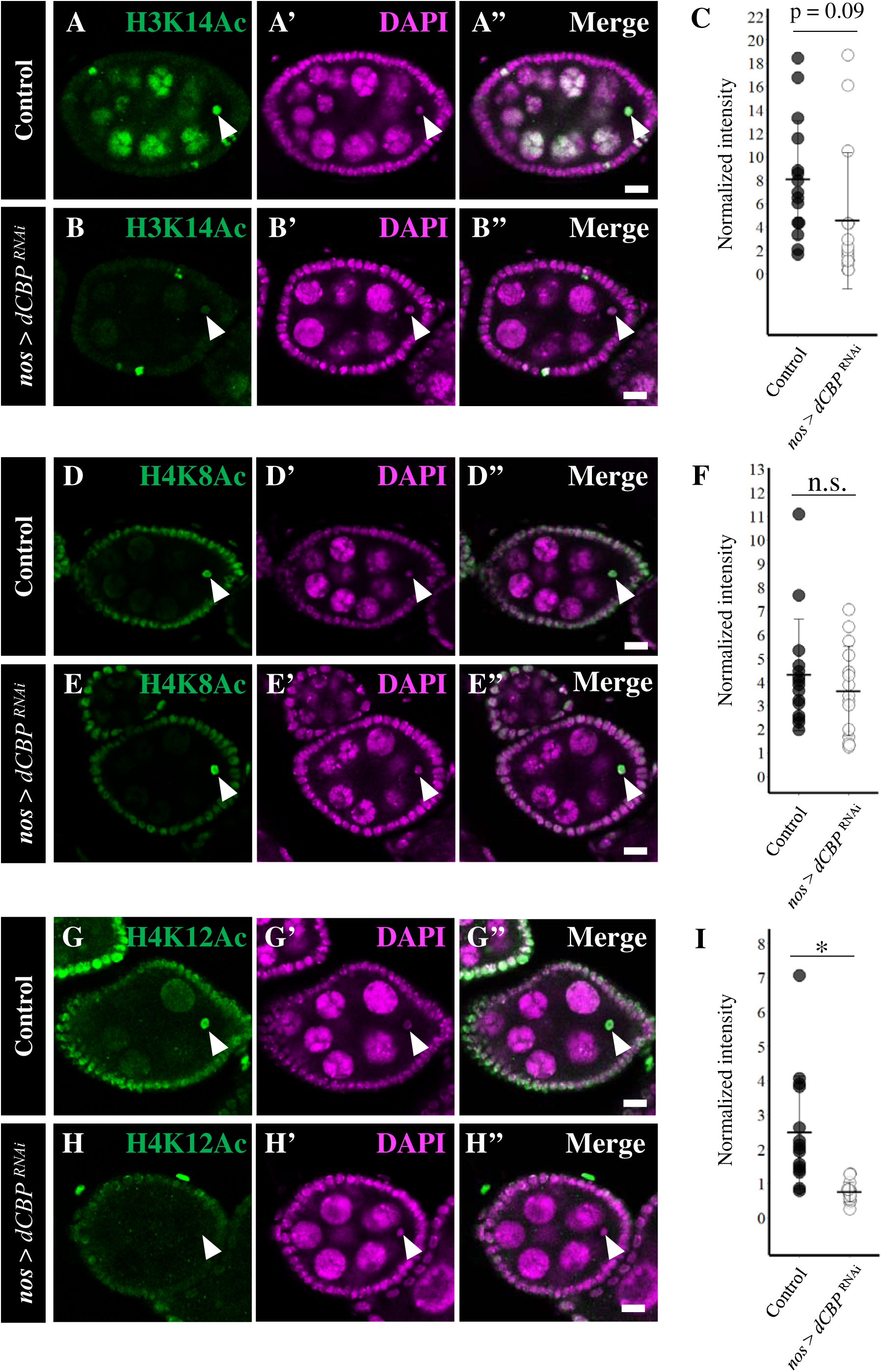
Effects of *dCBP* germline-specific knockdown on histone lysine acetylation in the karyosome. (A–C) H3K14 acetylation in control (A–A’’) and germline-specific *dCBP*-knockdown (B–B’’) karyosomes. Quantification of normalized karyosome signal intensity is shown in (C) (control, 8.05±5.05 S.D.; *nos* > *dCBP*^RNAi^, 4.53±5.81 S.D.; p =0.09, Welch’s t-test). (D–F) H4K8 acetylation in control (D–D’’) and germline-specific *dCBP*-knockdown (E–E’’) karyosomes. Quantification of normalized karyosome signal intensity is shown in (F) (control, 4.31±2.35 S.D.; *nos* > *dCBP*^RNAi^, 3.61±1.89 S.D.; p > 0.1, Welch’s t-test). (G–I) H4K12 acetylation in control (G–G’’) and germline-specific *dCBP*-knockdown (H–H’’) karyosomes. Quantification of normalized karyosome signal intensity is shown in (I) (control, 2.48±1.64 S.D.; *nos* > *dCBP*^RNAi^, 0.75±0.28 S.D.; p <0.05, Welch’s t-test). For normalization methods, see Materials and Methods. In all images, green indicates histone acetylation and magenta indicates DAPI staining of DNA. Arrowheads indicate karyosomes. Scale bars: 10 µm. Asterisks indicate p < 0.05, and n.s. indicates p > 0.1 (Welch’s t-test).

### dCBP regulates meiotic chromosomes and fertility

Our previous study showed that histone lactylation primarily occurs in the fully formed karyosome at stage 2 and diminishes as the karyosome begins to decondense during late oogenesis around stage 10 (Hayashi et al., 2025c). This time window overlaps with the period between maturation of the synaptonemal complex (SC), a key meiotic chromosome structure, and the onset of its disassembly. We therefore hypothesized that histone lactylation may contribute to SC maintenance and examined whether *dCBP* knockdown affects SC integrity.

To this end, we analyzed the localization of C(3)G, a transverse filament protein of the SC that is essential for SC formation and maintenance (Anderson et al., 2005, Page and Hawley, 2001, Hughes et al., 2018). In control ovaries, C(3)G accumulated on oocyte chromosomes by region 3 of the germarium (Figures 4A, A’) and was maintained in oocyte nuclei during early oogenesis (Figures 4B, B’). C(3)G was then gradually disassembled from oocyte chromosomes during mid-oogenesis (Figures 4C, C’). In *dCBP* knockdown oocytes, C(3)G accumulation on oocyte chromosomes occurred normally in region 3, suggesting that initial oocyte specification was not affected (Figures 4D, D’, and H). By contrast, C(3)G signals on oocyte chromosomes were precociously fragmented (Figure 4E), lost (Figure 4F), or disassembled (Figure 4G) during early oogenesis, indicating defective SC maintenance in *dCBP*-knockdown oocytes (Figure 4H, Supp. Table 2). These results suggest that *dCBP* is required for the maintenance of meiotic chromosome structure.

**Figure 4.**
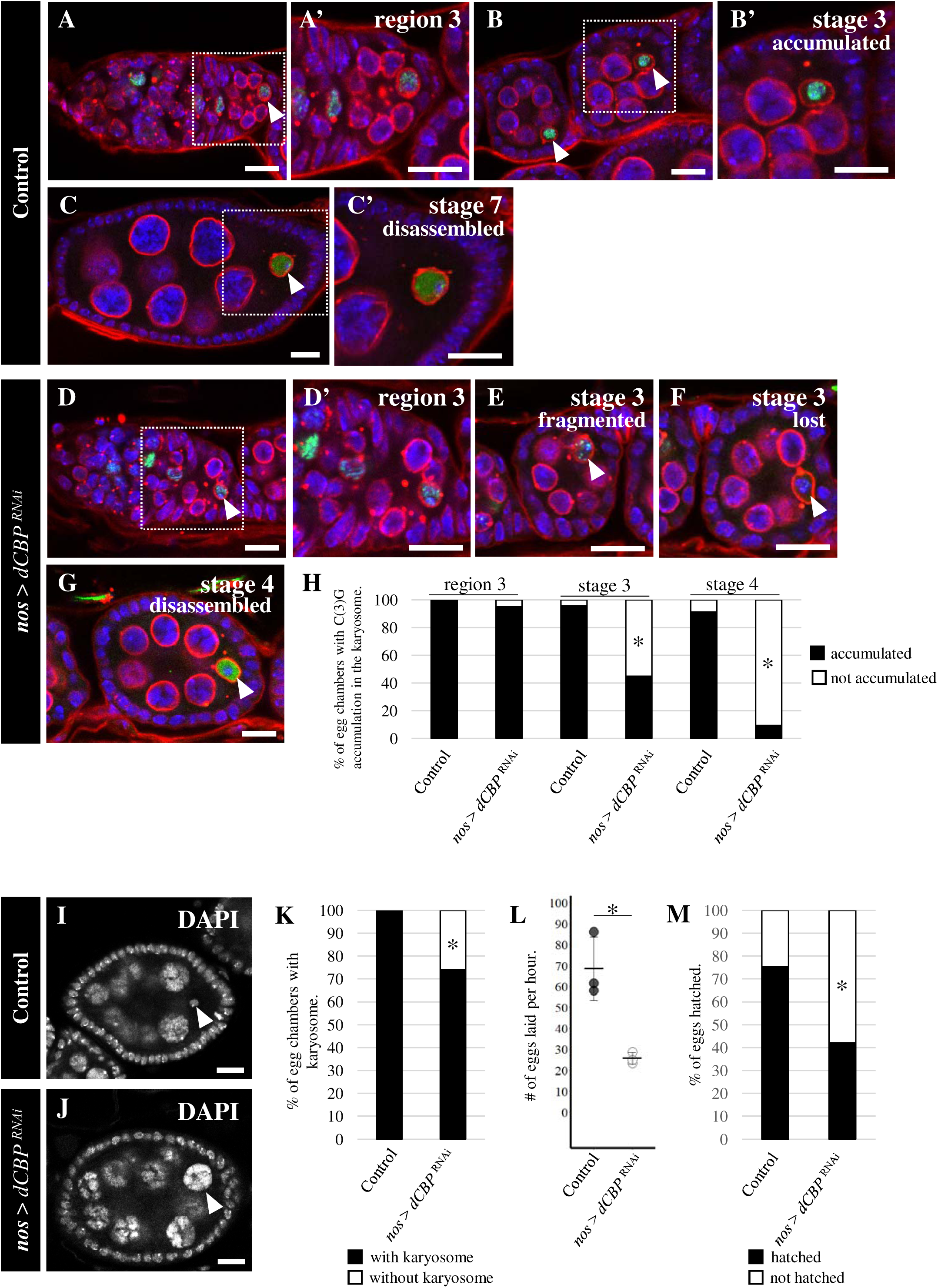
Effects of germline-specific *dCBP* knockdown on meiotic chromosomes and reproduction. (A–H) C(3)G localization in control (A–C’) and germline-specific *dCBP*-knockdown (D–G’) oocytes. C(3)G is shown in green, wheat germ agglutinin (WGA)-labeled nuclear membranes are shown in red, and DNA (DAPI) is shown in blue. Dotted squares indicate the regions enlarged in the corresponding magnified images (A’, B’, C’, D’, E’, F’, and G’). Statistical analysis is shown in (H). (I–K) Representative oocyte nuclear morphology in control (I) and germline-specific *dCBP*-knockdown (J) egg chambers. Statistical analysis is shown in (K). (L, M) Egg production (L) and embryonic hatching rate (M) in control and germline-specific *dCBP*-knockdown females. Scale bars: 10µm, Asterisks indicate p < 0.05 (H, K and M: Fischer’s exact test, L: Welch’s t-test).

Classical studies have shown that defects in SC maintenance impair oocyte fate maintenance and can cause oocytes to differentiate into nurse cells, resulting in the 16-nurse cell phenotype (Page and Hawley, 2001). Consistent with this, we observed a significant increase in the 16-nurse cell phenotype in *dCBP*-knockdown ovaries (Figures 4I–K, Supp. Movie 1, Supp. Table 3). We also observed reduced egg production in *dCBP*-knockdown females (Figure 4L). Furthermore, the hatching rate of embryos produced by *dCBP*-knockdown females was markedly decreased (Figure 4M, Supp. Table 4), suggesting that meiotic progression and/or subsequent embryonic development is impaired. Taken together, these observations raise the possibility that *dCBP*-mediated regulation of histone lactylation in the karyosome contributes to proper meiotic progression and female fertility.

Meiosis is a fundamental process underlying sexual reproduction and genomic diversity. During meiosis, chromosomes undergo extensive reorganization, including homolog pairing, synapsis, meiotic recombination, and large-scale chromatin remodeling. PTMs are recognized as crucial regulators of meiotic chromatin dynamics; however, most previous studies have focused primarily on repressive histone PTMs (Zhaunova et al., 2016, Navarro-Costa et al., 2016, Hu et al., 2022, Iovino et al., 2013, Feijao et al., 2022), reflecting the highly condensed and transcriptionally silent nature of meiotic chromosomes. By contrast, the contribution of histone acylation marks associated with active chromatin, such as histone lactylation, has remained largely unexplored. Here, our knockdown-based analyses provide evidence consistent with a model in which *dCBP*-mediated histone lactylation contributes to meiotic chromosome maintenance and normal oogenesis. To our knowledge, this study provides the first evidence implicating a CBP family protein in the regulation of histone lactylation within transcriptionally repressive meiotic chromatin.

CBP family proteins in mammals function as acyl-transferases involved in multiple histone acylation pathways, including acetylation and lactylation (Dancy and Cole, 2015, Zhang et al., 2019). In *Drosophila*, *dCBP* has also been implicated in histone acetylation, including H4K8 acetylation (Feller et al., 2015), although its involvement in histone lactylation has not previously been demonstrated. In this study, we found that *dCBP* knockdown preferentially affected H4K8 lactylation within the karyosome, whereas H4K8 acetylation was not uniquely enriched in this structure. Given that the karyosome represents a highly condensed and transcriptionally repressive meiotic chromatin domain, these observations raise the possibility that *dCBP*-mediated lactylation may play a more specialized role in meiotic chromatin regulation than acetylation during oogenesis. Such functions may extend beyond transcriptional regulation to include roles in higher-order chromatin organization and/or meiotic chromosome structure and stability.

The mechanism underlying the apparent selectivity of dCBP toward distinct acyl donors remains unclear. One possibility is that local metabolic states within oocytes influence the relative availability of acetyl-CoA and lactyl-CoA, both of which are ultimately derived from pyruvate metabolism. However, this interpretation may not fully explain our observations, as germline knockdown of *Ldh* did not markedly reduce karyosome lactylation. Alternatively, germline-specific cofactors may modulate *dCBP*-dependent lactylation in meiotic chromatin. In this context, the germline transcription factor *mamo*, which genetically interacts with *dCBP* and functions together with germline transcriptional factor *ovo* in germline gene regulation (Nakamura et al., 2019, Hayashi et al., 2025a, Alizada et al., 2025, Mukai et al., 2007), represents an intriguing candidate for future investigation. Although contributions from *dCBP*-dependent acetylation at other histone residues (Feller et al., 2015), as well as non-enzymatic functions of dCBP (Marsh et al., 2025, Hunt et al., 2022), cannot be excluded, the preferential reduction of lactylation together with the temporal coincidence between karyosome lactylation and SC maintenance is consistent with a model in which d*CBP*-mediated lactylation contributes to meiotic chromosome maintenance. Future studies identifying germline-specific cofactors of dCBP and their relationship to histone lactylation will help clarify the mechanistic basis of this regulation.

Another unexpected finding of this study was the relatively modest effect of germline *Ldh* knockdown on karyosome lactylation. One possible explanation is that the knockdown efficiency achieved by the RNAi line used here was insufficient, as this RNAi strain was reported to produce a substantially weaker reduction in lactate production than the corresponding deletion mutant (Li et al., 2019). Alternatively, lactate supplied from surrounding somatic cells may contribute to karyosome lactylation. Lactate is known to be exported through monocarboxylate transporters (MCTs)(Zhang et al., 2024), and recent studies in mammals demonstrated that MCT-dependent lactate transport contributes to CBP-dependent histone lactylation (Wang et al., 2022, Yang et al., 2022b). Notably, our previous work suggested that germline and somatic cells exhibit distinct metabolic states during oogenesis (Hayashi et al., 2025b), raising the possibility that metabolic interactions between these cell types influence meiotic chromatin regulation.

Future studies examining follicle cell-specific disruption of lactate production or transport will therefore be important to clarify whether somatically derived lactate contributes to karyosome lactylation and meiotic chromosome maintenance. If such perturbations phenocopy germline *dCBP* knockdown, this would further support a model in which lactate-dependent chromatin regulation contributes to meiotic chromosome maintenance.

Histone lactylation is an evolutionarily conserved histone PTM that links cellular metabolism to chromatin regulation (Meng et al., 2021, Zhang et al., 2019). Meiosis is also widely recognized as a metabolically sensitive process across diverse eukaryotes, and histone lactylation during meiosis has been observed in mammals (Borner et al., 2023, Zhang et al., 2021, Yang et al., 2022a, Hayashi et al., 2025c). Our findings therefore raise the possibility that metabolic regulation of meiotic chromatin through histone lactylation may represent a broadly conserved mechanism in animal reproduction. Further investigation of lactylation-mediated chromatin regulation in meiosis may help uncover how metabolic states directly influence chromosome dynamics and reproductive capacity.

### Study limitations

Although our study identifies *dCBP* as a regulator of histone lactylation in the karyosome, whether dCBP directly mediates histone lactylation in meiotic chromatin remains unclear. In addition, because *dCBP* depletion also partially affected histone acetylation, future studies will be required to distinguish the respective contributions of lactylation and acetylation to meiotic chromosome regulation.

## Materials & Methods

### Fly strains

Flies were maintained on standard *Drosophila* medium at 25 LJ. The wild-type strain used in this study is Oregon-R. For Gal4-based knockdown experiments, adult flies were maintained at 29 LJ for three days to enhance RNAi efficiency. Germline-specific Gal4 driver *nanos*-Gal4::VP16 (*nos*-Gal4, gift from D. Van Doren)(Van Doren et al., 1998) was crossed to VALIUM20-based UAS-RNAi strains [Strains from Bloomington Drosophila Stock Center (BDSC); UAS-*Ldh* RNAi (#33640), UAS-*AcCoAS* RNAi (#41917), UAS-*CG6432* RNAi (#44565), UAS-*Tip60* RNAi (#35243), UAS-*dCBP* RNAi (#37489), UAS-*AlaRS* RNAi (#43274), Strain from National Institute of Genetics (NIG) Japan; UAS-*Gcn5* RNAi (#HMS00941)]. A strain having an empty attP2 landing site (#36303, BDSC) was crossed to *nos*-Gal4 as a control for knockdown experiments.

### Immunohistochemistry

Immunostaining of *Drosophila* ovaries was performed according to standard procedures (Hayashi et al., 2025c). The ovaries of adult flies were dissected in Phosphate-Buffered Saline (PBS) and fixed with 4% Paraformaldehyde (PFA) in PBS for 15 minutes. We used 2% Bovine Serum Albumin (BSA), 0.1% Tween-20, 0.1% Triton X-100 in PBS as blocking solution and antibody diluent. The following primary antibodies were used: mouse anti-C(3)G antibody [1:500, kind gift from Drs Hughes and Billmyre (Anderson et al., 2005)]. The primary antibodies against lactylated lysine: rabbit anti-Lactyl-Lysine (Kla) antibody [PTM Biolabs (Chicago, IL, U.S.A.), PTM-1401, 1:500], anti-Lactyl-HistoneH3-Lysine14 (H3K14la) (PTM-1414, 1:2000), H4K8la (PTM-1415, 1:2000), H4K12la (PTM-1411, 1:2000). The secondary antibodies used were purchased from Thermo Fisher Scientific (Thermo Fisher)(Waltham, MA, U.S.A.): anti-rabbit IgG Alexa Fluor 488 plus (1:500, A32731), anti-mouse IgG Alexa Fluor 488 plus (1:500, A32723). Nucleic acid staining was performed with DAPI (1:500, D523, DOJINDO Laboratories, Kumamoto, Japan). Nuclear membrane staining was performed with Wheat Germ Agglutinin Alexa 568 (1:250, W56133, Thermo Fisher). Stained ovaries were mounted in ibidi mounting medium (ibidi, Grafelfing, Germany) and imaged basically using a confocal microscope (Stellaris8, Leica Microsystems, Wetzlar, Germany), except the counting of 16-nurse cell phenotype, in which we used a wide-field fluorescence microscope (DM6000, Leica Microsystems).

### Image quantification

Quantification of fluorescence intensity in the karyosome was performed using Fiji (ImageJ 1.54p). Images used for quantification were acquired at 12-bit depth, and detector gain was adjusted so that karyosome signals in mid-oogenesis were near saturation (maximum A.U.: 4096). To normalize intensity differences between samples, the average fluorescence intensity of three adjacent posterior follicle cell nuclei within the same focal plane was used as an internal reference. Regions of interest were defined based on DAPI staining. Statistical analyses were performed using R (version 4.5.0).

### Analysis of egg production and hatching rate

Twenty virgin females from control or dCBP-knockdown strains were crossed with 20 wild-type males for 2 days. Eggs were collected on egg-laying plates containing 50% grape juice, 2% agar, 1% ethanol, and 1% acetic acid, and the total number of eggs was counted. After counting, eggs were transferred to fresh plates and incubated at 25 LJ for 24 h. The hatching rate was then calculated based on the number of hatched larvae.

### Expression analysis of candidate genes

Previously published ovarian single-cell RNA-seq data [GSE136162; (Rust et al., 2020)] were used to analyze the expression of candidate genes. The dataset was processed using Seurat for clustering and visualization of feature plots (Seurat version 5.3.0; R version 4.5.0).

## Acknowledgement

We thank the Bloomington Drosophila Stock Center, Developmental Studies Hybridoma Bank, and National Institute of Genetics (NIG)-Fly for providing us with the materials. We also thank Drs Hughes and Billmyre for providing us with the C(3)G antibody. We also thank the Center for Advanced Technical and Educational Supports, Faculty of Agriculture, Kyushu University, for providing facilities. This work was supported in part by Grants-in-Aid for Scientific Research from Japan Society for Promotion of Science (JSPS) (KAKENHI Grant Number: 26K02041, 23K05777 to Y.H.)

## Author contributions

K. N., D.S., and Y. H. designed the experiments; K. N. performed the experiments; K. N. and Y. H. wrote the paper. All authors reviewed the manuscript.

---

**Supplemental Figure 1.**
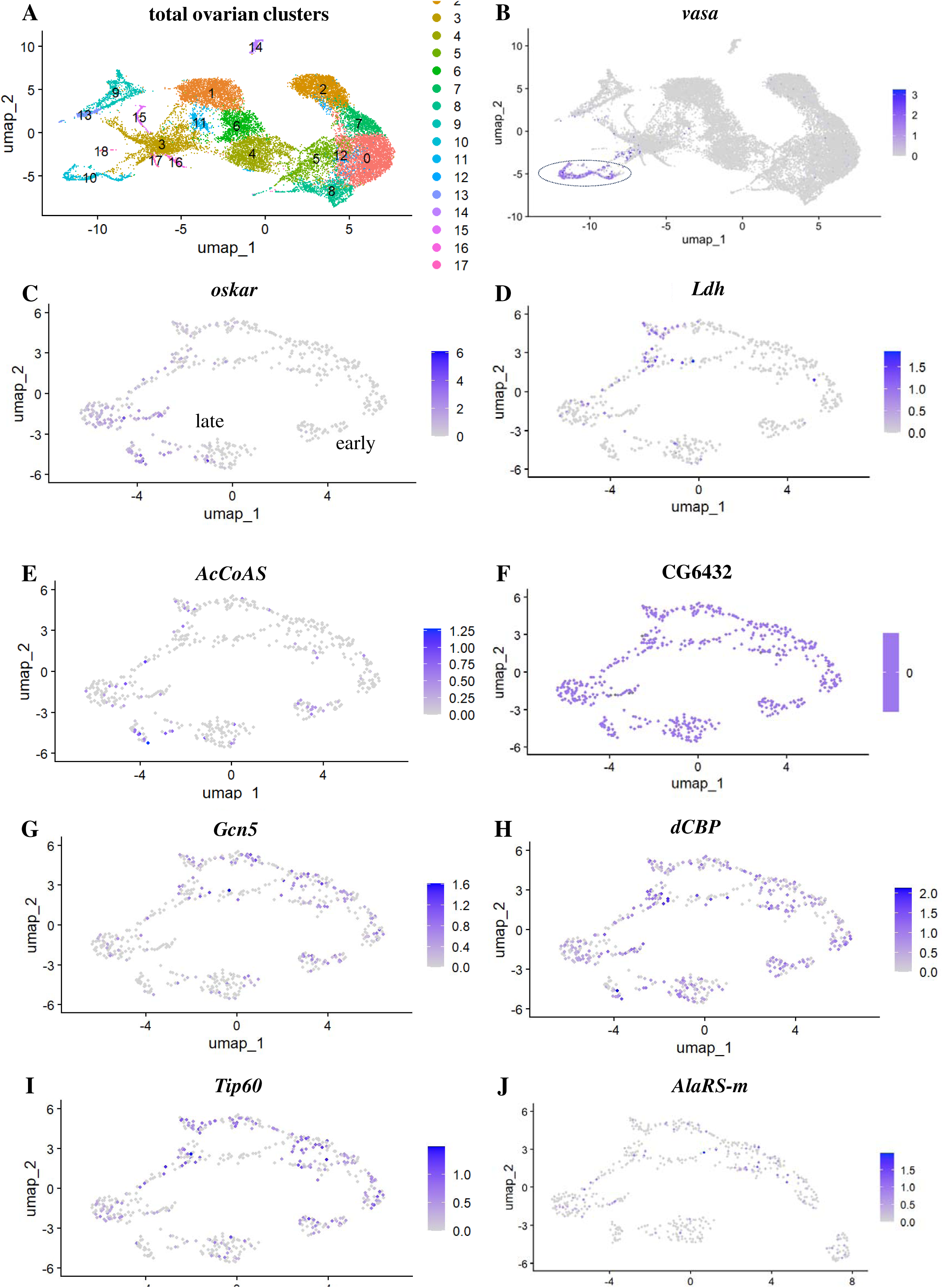
Expression of candidate genes in the ovarian germline. To analyze the expression of candidate genes, we used a published single-cell RNA-seq dataset of Drosophila ovarian cells (GSE136162).(A–C) Cluster 10 from the total cell clusters (A) was identified as the germline cluster based on expression of the germline marker *vasa* (dotted circle in B). Progression of germline development within the cluster was evaluated using the late germline marker *oskar* (C).(D–J) Expression patterns of candidate genes within the germline cluster (cluster 10). Gene names are indicated above each panel.

**Supplemental Figure 2.**
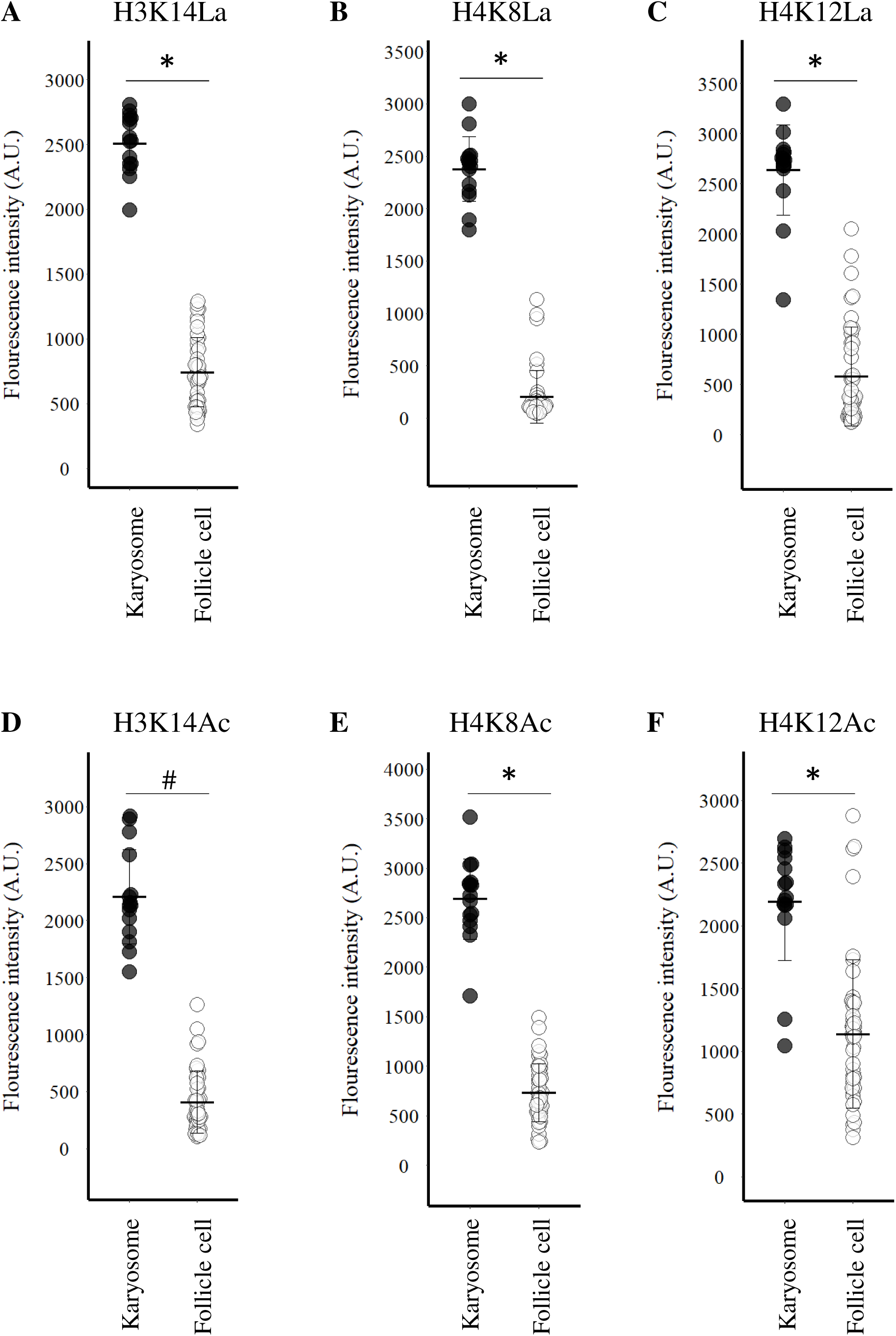
Quantification of histone lactylation and acetylation signals in karyosomes. (A–C) Quantification of H3K14La (A), H4K8La (B), and H4K12La (C) fluorescence intensity in karyosomes and follicle cells (H3K14La: karyosome, 2508.33±227.66 S.D.; follicle cell, 742.71±268.45 S.D.; p<0.05, Student’s t-test. H4K8La: karyosome, 2376.78± 308.58 S.D.; follicle cell, 202.64±248.19 S.D.; p<0.05, Student’s t-test. H4K12La: karyosome, 2635.74±447.48 S.D.; follicle cell, 580.22±490.62 S.D.; p<0.05, Student’s t-test). (D–F) Quantification of H3K14Ac (D), H4K8Ac (E), and H4K12Ac (F) fluorescence intensity in karyosomes and follicle cells (H3K14Ac: karyosome, 2209.54±415.02 S.D.; follicle cell, 407.10±273.75 S.D.; p<0.05, Welch’s t-test. H4K8Ac: karyosome, 2688.11±403.71 S.D.; follicle cell, 735.57±292.26 S.D.; p<0.05, Student’s t-test. H4K12Ac: karyosome, 2192.50±467.91 S.D.; follicle cell, 1137.52±593.82 S.D.; p<0.05, Student’s t-test). All images were acquired at 12-bit depth, with karyosome signals near saturation (4096 A.U.). Asterisks indicate p < 0.05 (Student’s t-test), and # indicates p < 0.05 (Welch’s t-test).

**Supplemental Table 1. Results of germline-specific knockdown of candidate genes.**

**Supplemental Table 2. C(3)G enrichment in oocyte nuclei.**

**Supplemental Table 3. Percentage of egg chambers with normally formed karyosomes.**

**Supplemental Table 4. Hatching rate of embryos laid by control and *dCBP*-knockdown females.**

**Supplemental Movie 1. Three-demensional view of the 16-nurse-cell phenotype in *dCBP*-knockdown ovaries**. Nuclei are visualized with DAPI. Scale bar: 10µm.

